# Establishment and imaging evaluation of Guangxi Ba-Ma mini-pigs as a model ofcerebral hernia induced by acute intracranial hypertension

**DOI:** 10.1101/487967

**Authors:** Jianguo Zhou, Lixuan Huang, Xiaoling Zhu, Wupeng Wei, Yongbiao Feng, Xiangfei Ma, Ling Zhang, Gang Zeng, Jianfeng Zhang, Weixiong Li, Huamin Tang

## Abstract

In order to establish an animal model of cerebral hernia induced by acute intracranial hypertension for subsequent research in emergency medicine. Six Guangxi Ba-Ma mini-pigs which were injected with autologous arterial blood by micropump to induce intracranial hematoma in the frontal and temporal parietal lobes. Changes in intracranial pressure (ICP), mean arterial blood pressure (MAP) and cerebral perfusion pressure (CPP)were observed during hematoma formation. Morphological changes and the occupying effects of intracranial hematoma were analyzed using computed tomography (CT) and magnetic resonance imaging (MRI) of the head. ICP, MAP and CPP gradually increased during blood injection, and they began to slowly decrease at 10 min after injection, though they remained higher than before injection. These three parameters were significantly different before blood injection, immediately after injection and at 10 min after injection (P< 0.05). Head CT and MRI showed cerebral hernia induced by acute intracranial hypertension. These results demonstrate micropump injection of autologous arterial blood can lead to acute intracranial hypertension in mini-pigs, which may be a useful model of cerebral hernia.

## INTRODUCTION

Many diseases can lead to increased intracranial pressure (ICP), such as traumatic brain injury, hypertensive cerebral hemorrhage, brain tumor, meningitis, and hepatic encephalopathy[10]. Some researchers have used rats, rabbits and pigs as experimental animals to study intracranial hypertension[14,12,13], but few studies have described animal models of brain hernia. Here we describe the Guangxi Ba-Ma mini-pigas a potential model of cerebral hernia induced by acute intracranial hypertension in order to lay a subsequent research in emergency medicine.

## METHODS

### Experimental animals

Experimental procedures were approved by the Animal Ethics Committee of The Second Affiliated Hospital of Guangxi Medical University.Six healthy male and female Guangxi Ba-Ma mini-pigs 7-8 months old were obtained from the College of Animal Science and Technology of Guangxi University. Mean body weight was 26.90±3.17 kg (range, 24-32 kg). Pigs were free to drink food and water before surgery in order to avoid hypoglycemia when they fast 5 h beforehand [11]. Animals were maintained at 25 °C.

### Analgesia and anesthesia

Animals were injected intramuscularly with ketamine hydrochloride (20 mg/kg). They were anesthetized via slow venous infusion of 10% chloral hydrate (Chengdu Chron Chemicals, Chengdu, China) into the ear vein (3-4 ml/kg). For subsequent surgical procedures, the animal was fixed on the operating table in a supine position. Lodophor and alcohol disinfection and sterile surgical towels were used.

### Tracheotomy and electrocardiography (ECG)

The skin on the chest was shaved, an electrode was placed over the heart, and 1% lidocaine was infused to provide local anesthesia. Skin and subcutaneous tissue were cut along the median line of the neck. The anterior cervical muscle group was separated, tracheotomy was performed on the 2-3 ring below the annular cartilage, and a 6.5-mm tracheal catheter with balloon (Smith Medical Devices, UK) was implanted. The skin was sutured, and the tracheal catheter was fixed.

### Mean arterial blood pressure (MAP), intracranial pressure (ICP) and cerebral perfusion pressure (CPP) monitoring

The groin was disinfected, sterile surgical towels were placed, and an incision approximately 5 cm long was made along the inguinal ligament. Subcutaneous tissue, superficial fascia, deep fascia and muscles were separated layer by layer. Blood vessels and nerves were fully exposed (Figure 1), including the femoral vein (V) on the medial side, femoral artery (A) in the middle, and femoral nerve (N) on the outside (V-A-N order). The femoral artery was carefully separated. A sensor was implanted into the right femoral artery and connected to an external ECG monitor to continuously monitor heart rate, respiratory rate and MAP. An arterial catheter was inserted into the left femoral artery, and blood gases were analyzed. This catheter was also used to perform intracranial injection as described below.

**Figure 1.**
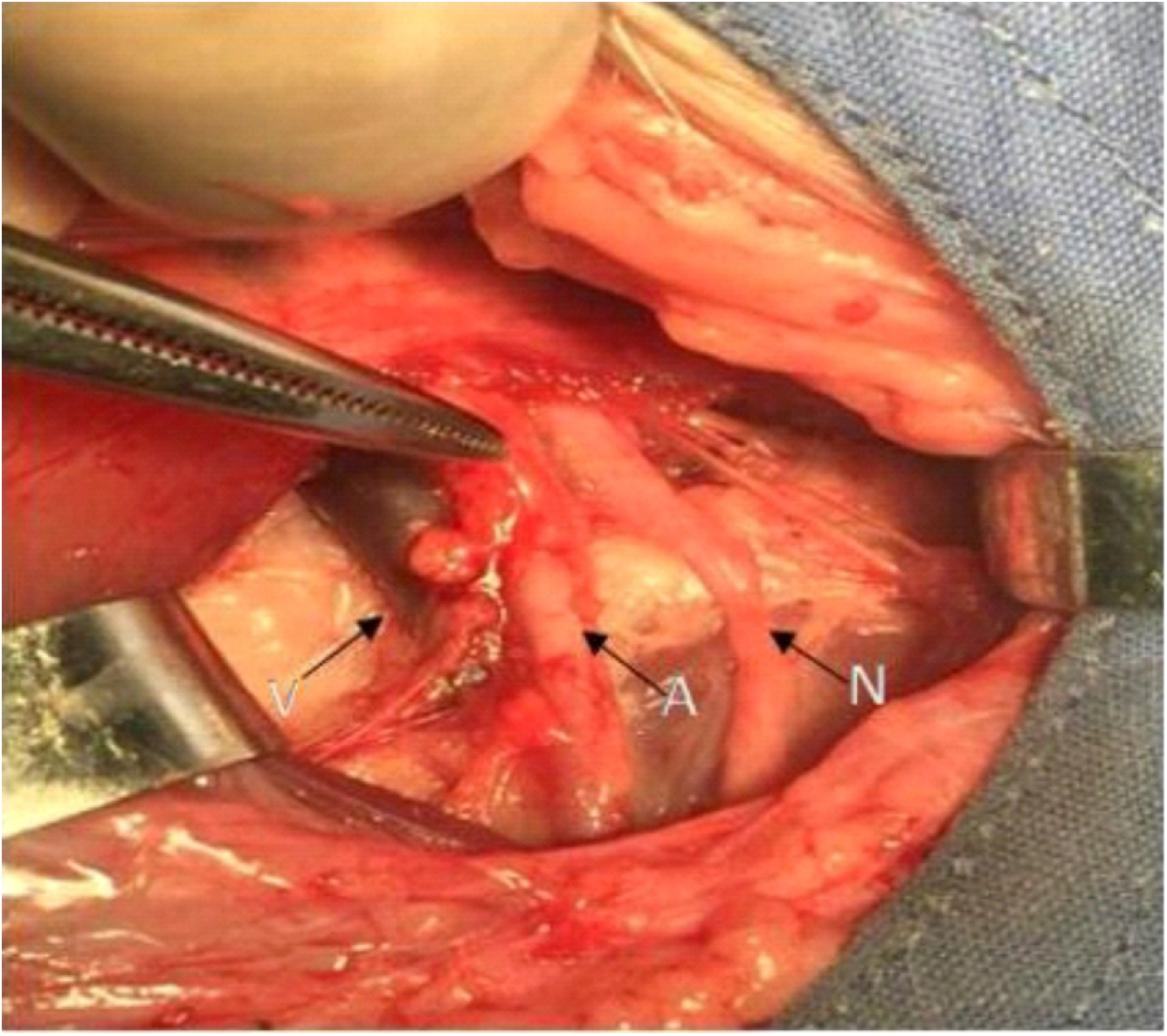
Anatomical location of the femoral arteries (*left* to *right*): femoral vein (V), femoral artery (A) femoral nerve (N).

The skin on the head was shaved and disinfected, and sterile surgical towels were spread. The scalp was cut in a "+" shape with a diameter of 2 cm, and the skull was drilled with a diameter of 0.8 cm at a location 1 cm before the front of the binaural line and 1.5 cm on the left side of the sagittal seams (Figure 2). The same method was used on the right side of the head to continuously monitor ICP (Sophysa, France). The ICP optical fiber probe was zeroed, then placed into the brain parenchyma from the back of the right bone foramen. The probe was connected to an external monitor. CPP was calculated from MAP and ICP data using the formula [5]: CPP (mmHg) = MAP (mmHg) - ICP (mmHg).

**Figure 2.**
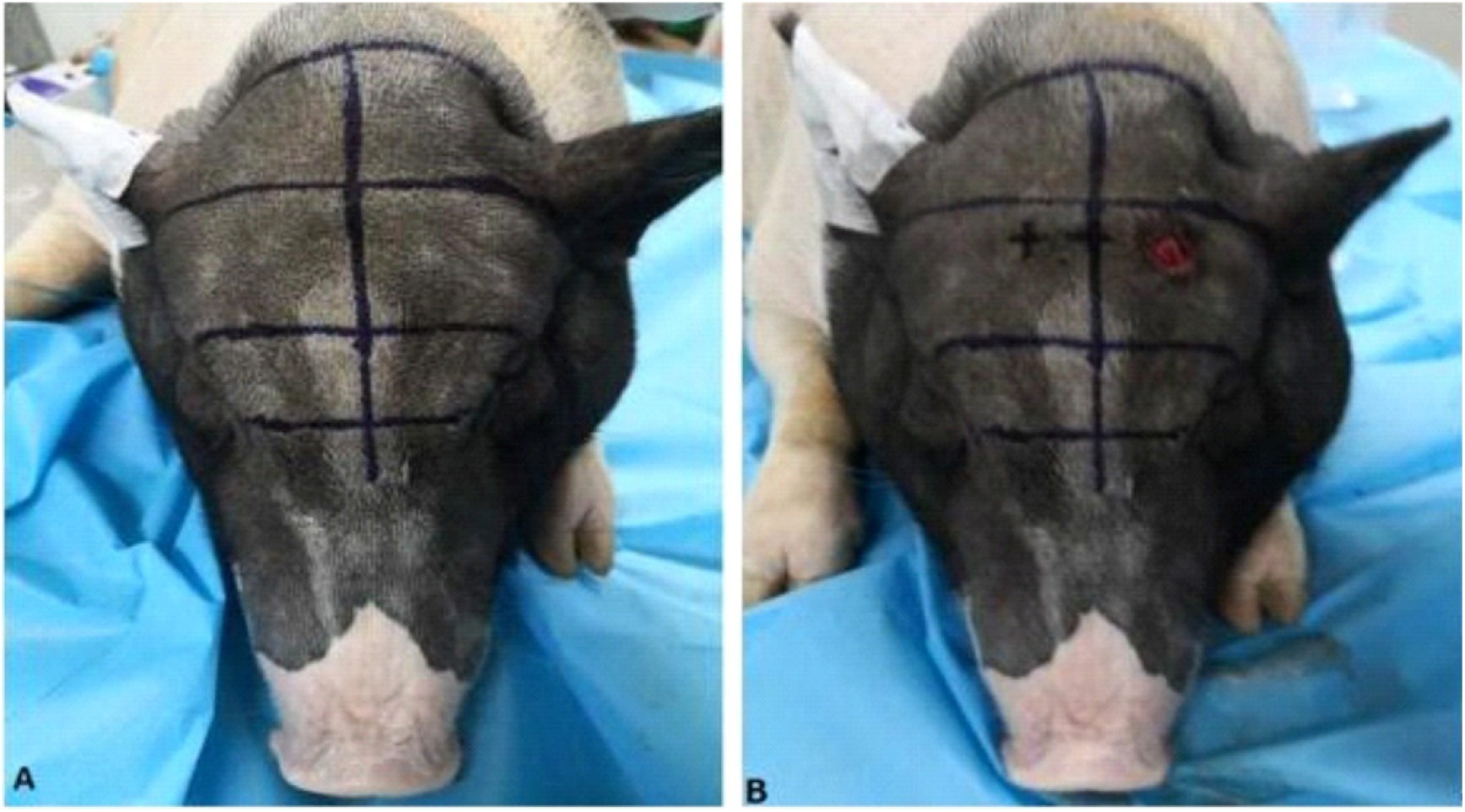
Positional maps of the Guangxi Ba-Ma mini-pig skull (A) before and (B) after drilling.

### Induction of cerebral hernia by acute intracranial hypertension

In the left bone foramen, the detaining trocar was used to puncture the dura mater about 0.5 cm vertically without cutting. The metal core was removed and the bone foramen was closed with bone wax. After heparinization with a 20-ml syringe, left femoral artery blood was drawn. Arterial blood was injected at 1 ml/min into the frontal and temporal parietal lobes with a micropump through a catheter-connected puncture needle in order to induce acute intracranial hypertension. ICP, MAP, heart rate, respiratory rate and pupil diameter were continuously monitoredduring and after blood injection. When injection of autologous arterial blood was complete, the bone foramen was quickly closed with bone wax.

### Computed tomography (CT) and magnetic resonance imaging (MRI) of the head

After injection of arterial blood, morphological changes and occupying effects of the intracranial hematoma induced by acute intracranial hypertension were analyzed using 128-slice dual-source spiral CT (Siemens, Germany) at a layer thickness of 5 mm, and using a 3.0-T MRI (GE, USA) at a layer thickness of 4 mm.

## Statistical analysis

Data were analyzed using SPSS 20.0 (IBM, USA), and data were expressed as mean ± standard deviation. ICP, MAP and CPP were compared before blood injection, immediately after injection and 10 min after injection using the paired *t* test. Significance was defined as *P*< 0.05.

## RESULTS

Acute intracranial hypertension and subsequent cerebral hernia were established successfully in all six Guangxi Ba-Ma mini-pigs without any death. Volume of intracranial hematoma ranged from 11.50 to 25.00 ml (mean, 16.75±4.95 ml).

### Changes in ICP, MAP and CPP

Before blood injection, ICP ranged from 5 to 12 mmHg (mean, 8.2±2.5 mmHg). ICP gradually increased during injection; the highest value was 51.3±2.6 mmHg. At the end of injection, ICP slowly decreased. At 10 min after injection, the mean value was 39.8±2.1 mmHg, which was higher than pre-injection (Table 1). ICP differed significantly before injection, immediately after injection and 10 min after injection (*P* < 0.05).

**Table 1.** Comparison of ICP before autologous arterial blood injection, immediately after injection and 10 min after injection

| Animal | ICP (mmHg) |  |  |
| --- | --- | --- | --- |
|  | Before injection | Immediately after injection | 10 min after injection |
| 1 | 12 | 49 | 42 |
| 2 | 9 | 54 | 37 |
| 3 | 8 | 49 | 42 |
| 4 | 5 | 55 | 38 |
| 5 | 9 | 51 | 39 |
| 6 | 6 | 50 | 41 |
| Mean | 8.2±2.5 | 51.3±2.6 | 39.8±2.1 |

Pre-injection vs. immediately after injection: $t = 23.545$ , $P < 0.001$
Pre-injection vs. 10 min after injection: $t = 28.387$ , $P < 0.001$
Immediately after injection vs. 10 min after injection: $t = 6.191$ , $P = 0.002$ .

Mean MAP before blood injection was 84.2±10.1 mmHg. MAP gradually increased during injection; the highest value was 179.7±6.3 mmHg. At the end of injection, MAP decreased slowly, and the mean value at 10 min after completion was 143.7±7.8 mmHg, which was higher than pre-injection (Table 2). MAP differed significantly before injection, immediately after injection and 10 min after injection (*P* < 0.05).

**Table 2.** Comparison of MAP before autologous arterial blood injection, immediately after injection and 10 min after injection

| Animal | MAP (mmHg) |  |  |
| --- | --- | --- | --- |
|  | Before injection | Immediately after injection | 10 min after injection |
| 1 | 89 | 176 | 149 |
| 2 | 78 | 177 | 151 |
| 3 | 99 | 171 | 150 |
| 4 | 73 | 181 | 141 |
| 5 | 90 | 185 | 131 |
| 6 | 76 | 188 | 140 |
| Mean | 84.2±10.1 | 179.7±6.3 | 143.7±7.8 |

Pre-injection vs. immediately after injection: $t = 16.025$ , $P < 0.001$
Pre-injection vs. 10 min after injection: $t = 12.411$ , $P < 0.001$
Immediately after injection vs. 10 min after injection: $t = 6.609$ , $P = 0.001$

Mean CPP before blood injection was 76.0±8.9 mmHg. CPP gradually increased during injection; the highest value was 128.3±6.3 mmHg. Immediately after injection, CPP decreased slowly, falling to a mean value of 103.8±7.7 mmHg by 10 min after injection, which was still higher than pre-injection (Table 3). CCP differed significantly before injection, immediately after injection and 10 min after injection (*P* < 0.05).

**Table 3.** Comparison of CCP before autologous arterial blood injection, immediately after injection and 10 min after injection

| Animal | CCP (mmHg) |  |  |
| --- | --- | --- | --- |
|  | Before injection | Immediately after injection | 10 min after injection |
| 1 | 77 | 127 | 107 |
| 2 | 69 | 123 | 114 |
| 3 | 91 | 122 | 108 |
| 4 | 68 | 126 | 103 |
| 5 | 81 | 134 | 92 |
| 6 | 70 | 138 | 99 |
| Mean | 76.0±8.9 | 128.3±6.3 | 103.8±7.7 |

Pre-injection vs. immediately after injection: $t = 10.528$ , $P < 0.001$
Pre-injection vs. 10 min after injection: $t = 5.556$ , $P = 0.003$
Immediately after injection vs. 10 min after injection: $t = 4.499$ , $P = 0.006$ .

### Head CT and MRI scan

Head CT revealed that the hematoma was located in the parenchyma of frontal temporal parietal lobe (Figure 3). T2-weighted MRI showed that the prepontine cistern and sulcus existed as a line-like high signal before injection and disappeared after injection (Figure 4). These results suggest that the cerebral hernia displaced brain tissue and compressed the brain stem.

**Figure 3.**
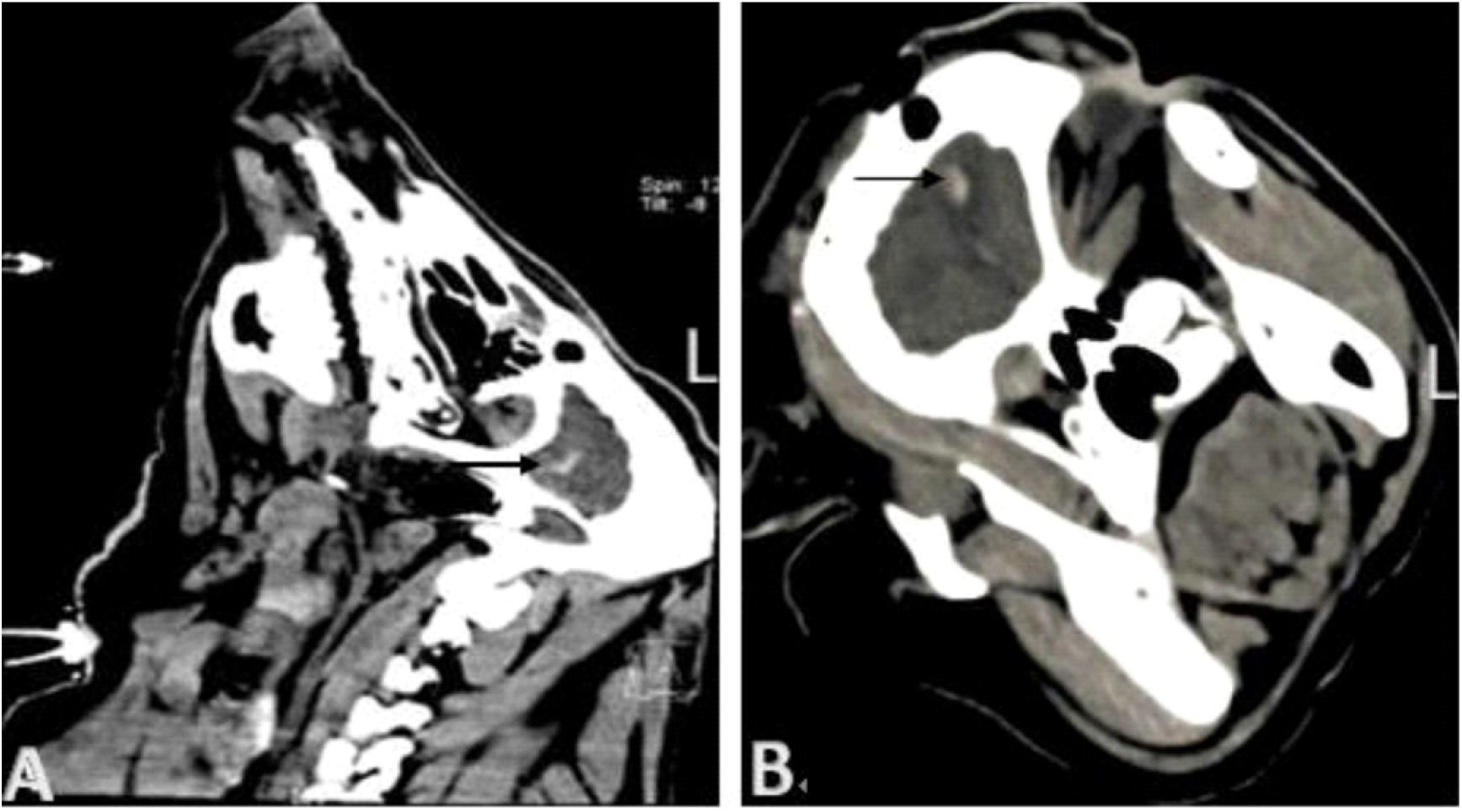
Head CT showing intracranial hematoma (black arrow) on the left side in (A) sagittal section and (B) coronal section.

**Figure 4.**
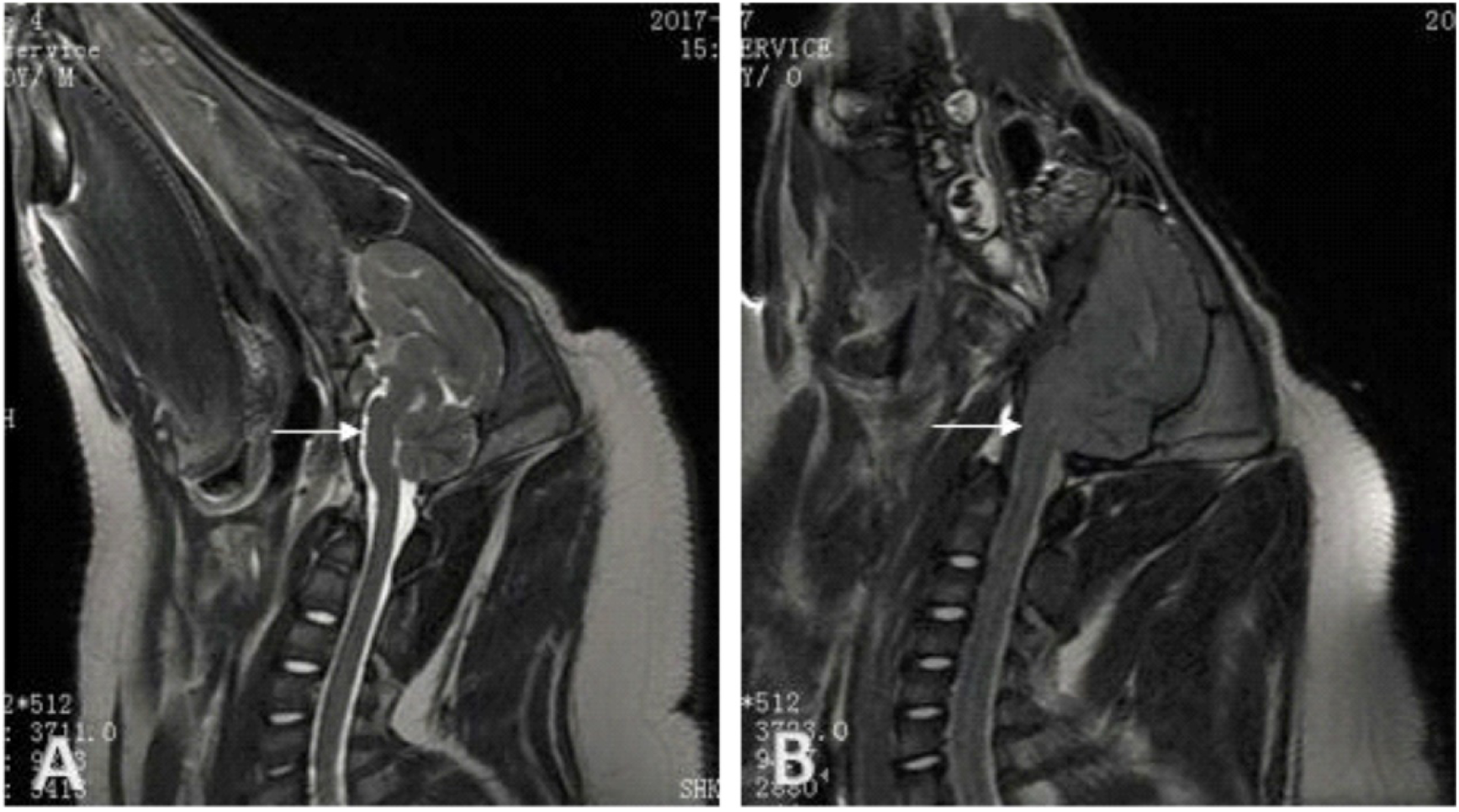
T2-weighted head MRI performed (A) before injection and (B) after injection. Before injection, the prepontine cistern and sulcus are visible as a line-like high signal (white arrow). Neither feature is visible after injection.

## DISCUSSION

Here we injected Guangxi Ba-Ma mini-pigs with hematoma in the frontal and temporal parietal lobe, and successfully established a stable model of cerebral hernia induced by acute intracranial hypertension. Guangxi Ba-Ma mini-pigs are intensively inbred for small body shape, well-defined biological characteristics and genetic stability, making them well-suited for experimental application. Their organ systems are similar to the human systems, and their hemodynamics are relatively stable.

Their craniocerebral volume is larger than that of rats and rabbits, and they can be injected with hematomas 20-30 timeslarger than rats. Their head can be analyzed easily using CT and MRI to observe hematoma, edema and changes in brain tissue metabolites[11]. Here we demonstrated that mini-pigs undergo cerebral hernia-related changes similar to humans in response to acute intracranial hypertension.

In our mini-pigs, the fixed size of the cranial cavity meant that as the intracranial hematoma gradually increased, pressure on the brain volume increased as well, leading to gradual increase in ICP and MAP. This is similar to the case of humans, where the Monro-Kellie doctrine predicts that because of the limited brain volume, ICP continues to increase due to cerebral edema, leading to tonsillar, callosus gyrusand subtentorial hernia [4]. Elevated ICP is linked to deterioration of neurological function in patients with trumatic brain injury[5]. In our model, at 10 min after blood injection, ICP and MAP began to decrease gradually, though they remained higher than pre-injection. This likely reflects certain compensatory functions in the brain, such as changes in cerebrospinal fluid and cerebral blood flow, which counteract damage arising from increasing intracranial hematoma pressure.

CPP drives cerebral blood flow: at constant cerebrovascular resistance, CPP correlates positively with cerebral blood flow[3]. CPP can fall because of a decrease in MAP, an increase in ICP or both[3]. When CPP decreases, cerebral blood flow may no longer be sufficient for adequate cerebral perfusion and oxygenation[6]. The CPP considered adequate varies among patients[10], and while consensus standards have not been defined, 50-60 mmHg is widely considered the minimum cerebral perfusion pressure required to prevent further brain injury[7]. In our mini-pig model, CPP increased during acute intracranial hypertension as a compensatory mechanism to ensure brain blood supply. The mini-pig model system may be useful for determining the most clinically relevant endpoints for predicting prognosis. Some researchers[9,1] believe that maintaining mean CPP as close as possible to the optimal cerebral perfusion pressure is key to favorable outcomes. Others advocate increasing MAP in order to maintain sufficient cerebral blood flow[8]. The Lund concept, in contrast, advocates reducing intravascular resistance and hydrostatic pressure and reducing cerebral blood volume in order to increase cerebral blood flow, despite lower CPP[2].

Our animals satisfied standard diagnostic criteria for acute intracranial hypertension (cerebral hernia) in mini-pigs, which means presenting at least three of the following symptoms: (1) ICP >50 mmHg, (2) MAP >150 mmHg, (3) slowed respiratory frequency (<6 beats/min) or tidal breathing or cessation, (4) heart rate <50 beats/min or >200 beats/min or presence of arrhythmia, (5) unilateral or bilateral pupil dilation to the margin, or (6) midline shift in intracranial hematoma or disappearance of prepontine cistern by head CT or MRI. Our research team chose to inject blood into the frontal and temporal parietal lobes in order to simulate trauma resulting, for example, from a high-altitude fall or blunt trauma. Frontal and temporal parietal lobe hemorrhage invades the cerebral ventricle and causes cerebral hernia, which quickly leads to respiratory depression, respiratory arrest, and sudden cardiac arrest. This type of injury shows abrupt onset and is associated with high rates of death and disability. Our mini-pig model may significantly advance research into emergency treatment of brain hernia.Our results with a small number of animals justify larger studies to explore factors associated with prognosis and to examine potential treatments.

## Acknowledgements

I sincerely thank Doctor Zhong Jian-Hong for his hard work and the revision of this article by Doctor A. Chapin Rodríguez.

## Compliance with ethical standards

### Ethical approval

All applicable international, national, and/or institutional guidelines for the care and use of animals were followed.

## Abbreviations and acronyms

CPP: cerebral perfusion pressure
CT: computed tomography
ECG: electrocardiography
ICP: intracranial pressure
MAP: mean arterial blood pressure
MRI: magnetic resonance imaging

